# Convergent neural dynamical systems for task control in artificial networks and human brains

**DOI:** 10.1101/2024.09.29.615736

**Authors:** Harrison Ritz, Aditi Jha, Nathaniel D. Daw, Jonathan D. Cohen

**Affiliations:** Princeton Neuroscience Institute, Princeton University, USA; Department of Statistics, Stanford University, USA; Department of Psychology, Princeton University, USA

**Keywords:** task switching, recurrent neural networks, state-space models, electroencephalography

## Abstract

The ability to switch between tasks is a core component of human intelligence, yet a mechanistic understanding of this capacity has remained elusive. Long-standing debates over how task switching is influenced by preparation for upcoming tasks or interference from previous tasks have been difficult to resolve without quantitative neural predictions. We advance this debate by using state-space modeling to directly compare the latent task dynamics in task-optimized recurrent neural networks and human electroencephalographic recordings. Over the inter-trial interval, both networks and brains converged into a neutral task state, a novel control strategy that reconciles the role of preparation and interference in task switching. These findings provide a quantitative account of cognitive flexibility and a promising paradigm for bridging artificial and biological neural networks.

---

Humans have a remarkable capacity to flexibly adapt how they perform tasks [1–4]. This flexibility reflects a core feature of goal-directed cognition [5] and is a strong indicator of cognitive changes both across the lifespan [6, 7] and in mental health [8, 9]. Despite the centrality of cognitive flexibility in our mental life, we still have a limited mechanistic understanding of how people rapidly configure task processing to achieve their moment-to-moment goals.

There is broad agreement in the task-switching literature that two core factors influence the ability to switch rapidly between stimulus-response mappings, often termed ‘task sets’ [10, 11]. The first factor is ‘task set reconfiguration:’ when a new task is cued, using cognitive control to actively switch between task sets [12, 13]. The second factor is the influence of prior task states, originally cast in terms of ‘task set inertia:’ proactive interference from the previous task set that passively decays over time [14, 15]. There have been long-standing debates over the relative contributions of reconfiguration and inertia [2, 3], with behavioral and neuroimaging experiments finding support for either or both factors [16–21]. However, this theorizing at the level of ‘factors,’ rather than quantitative process models, has made it difficult to generate precise predictions that could help adjudicate between theories of task switching from brain imaging experiments.

Dynamical systems models, often cast in the form of recurrent neural networks (RNNs), offer a promising avenue for formalizing the influence of reconfiguration and prior states on task switching [7, 22–24]. In these models, reconfiguration arises from cue-dependent activity or gating, whereas the influence of prior state arises from recurrent activity that causes task representations to persist across trials. Despite the formal rigor of these models, they remain underexplored for characterizing how neural dynamics depend on prior states and reconfiguration [25, 26]. These models have often relied on hand-crafted mechanisms to implement different components of task switching, which may not map well onto distributed computations in the brain. Moreover, previous work has made limited attempts to directly link the rich representational dynamics in artificial neural networks to recordings of brain activity [25].

In the work reported here, we bridged between dynamical theories of cognitive control and their neural mechanisms by leveraging two key innovations. First, we developed a gated RNN model of task switching that used theory-motivated training curricula to isolate the effects of prior state and reconfiguration. Second, we developed an analysis pipeline for fitting large-scale dynamical systems to both the RNN model and two human electroencephalography (EEG) datasets, providing a common format to quantify and compare task-switching computations across modalities. Our findings identify a novel task switching strategy in RNNs: convergence towards a ‘neutral’ task state during the inter-trial interval (ITI), but only in neural networks trained to switch between tasks. Critically, we find that this strategy was closely mirrored in the EEG datasets, reconciling the influences of active reconfiguration and previous task state.

## Task-optimized RNNs learn to switch tasks

We used RNNs with gated recurrent units (GRU; [27]) to model how people flexibly switch between tasks (Method 1). These models have both the recurrent dynamics and gating mechanisms that have been proposed as core components of task switching, allowing them to ‘gate-in’ new task states [22–24]. Critically, these models provide us with full access to the latent computations. To isolate the effects of reconfiguration and prior tasks, we trained identical RNNs under the influence of two different task factors: ‘switch-training’ and ‘trial-spacing’ (Fig. 1A).

**Fig. 1:**
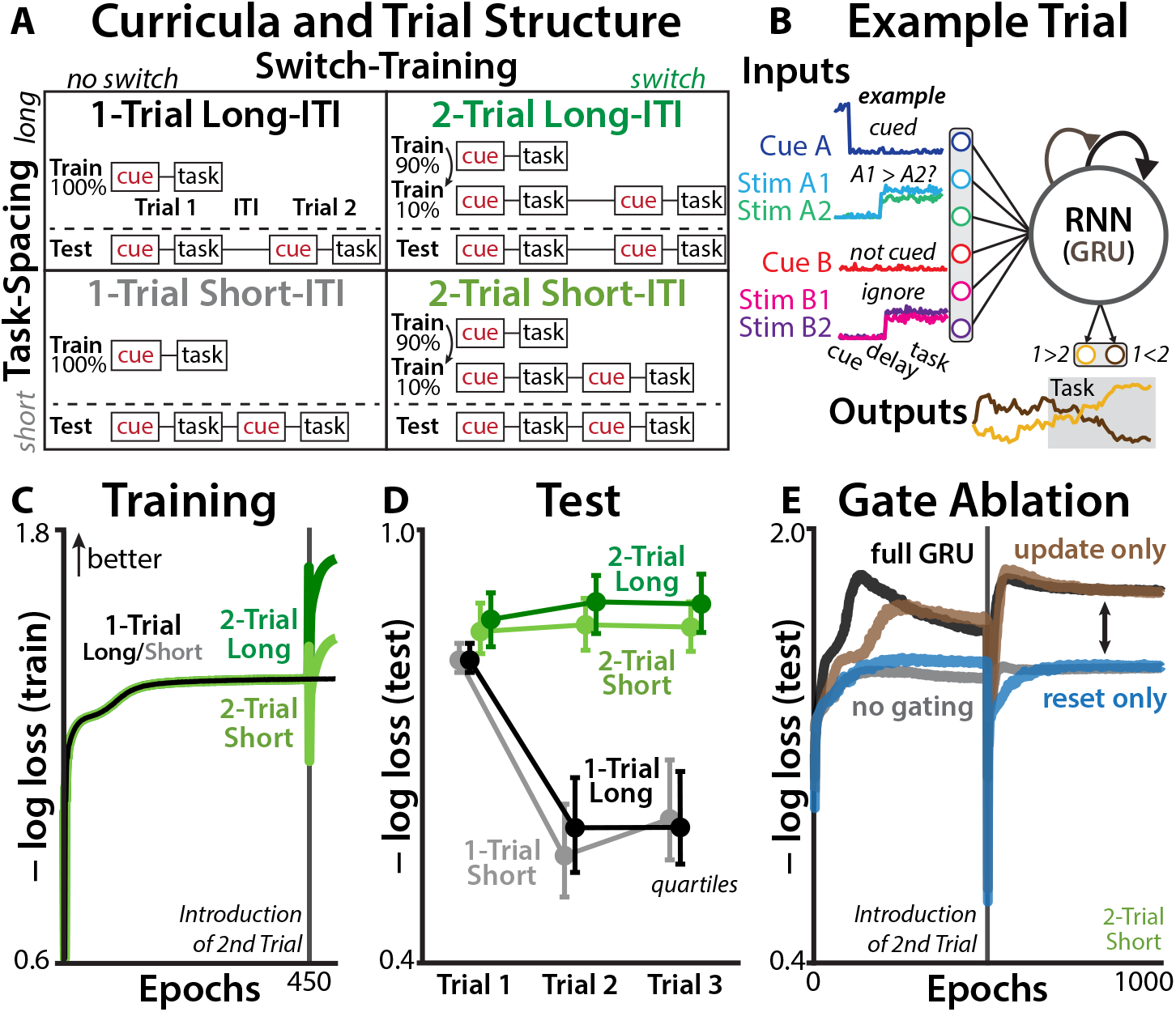
Training protocol for RNN models. **(A)** The *Switch-Training* curriculum (columns) manipulated RNNs’ experience with task-switching, capturing preparatory strategies. The *Task-Spacing* curriculum (rows) manipulated the inter-trial interval (ITI), capturing the influence of previous task. **(B)** Each trial consisted of a task cue (1-hot vector), a delay period (no inputs), and a target stimulus (two inputs per task). RNNs were trained to report which to the two task-relevant inputs had higher magnitude. **(C)** Training loss over training epochs. All networks had the same single-trial training for the first 450 epochs, and then 2-Trial networks were trained on pairs of trials. Input noise was included on training trials. **(D)** Differences in loss during the test phase (novel three-trial sequences, without input noise). Error bars reflect quartiles of the RNN distribution. **(E)** Test performance for RNNs with the full GRU architecture, either the update or reset gates fixed open, or no gating (‘Elman’ RNN). Shown for ‘2-Trial Short’ models on two-trial test sequences (no input noise).

The switch-training factor examined the role of learned control policies by varying the RNNs’ experience with switching between tasks. ‘1-Trial’ RNNs were trained to perform only one trial per sequence, which did not provide experience with task switching during learning. In each sequence, they received a one-hot task cue, a delay period, and then during a ‘task’ period they had to compare the relative magnitude of two task-relevant inputs (four simultaneous inputs, two per task; Fig. 1B; [28, 29]). RNN outputs were read out through a linear transformation and logistic nonlinearity, indicating the confidence for each response. We included a high level of input noise (cue SNR: 0 dB; trial SNR: -16.5 dB) to encourage evidence accumulation and top-down control.

Whereas 1-Trial RNNs only ever performed one trial per sequence, ‘2-Trial’ RNNs were also trained to switch between tasks (Fig. 1A, lower). 2-Trial RNNs performed single-trial sequences for the first 90% of their training, and then for the remaining 10% were trained with two trials per sequence (equal probability of switching or repeating tasks). This addition of task switching only late in training was designed to allow the network first to develop baseline competence in evidence accumulation, reproducing the emergence of control policies that accommodate over-learned task environments [30].

The ‘trial-spacing’ factor examined the influence of prior states by varying the ITI. This factor replicates classic behavioral experiments designed to test for inertia [16], capturing how task switching is influenced by the temporal proximity to the previous trial. Networks were trained with either short ITIs (20 timesteps; Fig. 1A) or long ITIs (60 timesteps).

After training these four sets of RRNs (N=512 per curriculum; Fig. 1C), we tested their generalization on novel three-trial sequences. All RNNs had fairly similar first-trial performance, suggesting a similar capacity for basic task processing (Fig. 1D). However, 2-Trial RNNs had much better performance on the second trial of the sequence, largely due to better performance on switch trials (Fig. S1). Despite their switch training, 2-Trial RNNs still had worse performance at shorter ITIs (Trial 2: *d* = 0.47; Trial 3: *d* = 0.53), consistent with a persistent influence of previous task states. On the third trial of the sequence, novel for all RNNs, 2-Trial RNNs maintained their performance advantage, consistent with a generalizable reconfiguration strategy.

We verified that gating mechanisms were used for active reconfiguration, consistent with prior theories of cognitive control [31, 32], by training RNNs with ablated GRU gates (Method 1.4). We found that update gates were necessary and sufficient for learning the two-trial sequences (Fig. 1E), consistent with a role for gating in active reconfiguration. Together, these simulations indicate that RNNs trained to switch between tasks developed generalizable gating policies, the dynamics of which we analyze further below.

### Quantifying RNNs’ latent task dynamics

We next turned to characterizing how RNNs’ latent task dynamics depended on switching-training and task-spacing. One common method for quantifying the dynamics of RNNs is finding locally linear representations around the model’s fixed points (or ‘slow points’; [33]). However, finding fixed points in input-driven networks can be difficult, and this procedure does not allow for direct comparisons to neural recordings. Instead, we linearized the entire RNN trajectory by fitting linear-Gaussian state-space models (SSM; often ‘latent linear dynamical systems’) to RNNs’ hidden unit activity [34, 35]. SSMs have long been used in computational neuroscience to characterize latent variables responsible for dynamics in multiunit neural recordings [36–38]. Despite the nonlinear activation functions in real and artificial neurons [39], linear models often provide highly predictive models of multi-scale brain activity [34, 40, 41]. Critically, linear SSMs can be characterized using a suite of tools from dynamical systems theory and control theory [42, 43], providing an interpretable alternative to nonlinear SSMs like hidden Markov models [44–46] or RNNs [47, 48].

The SSM generative model consists of a set of latent factors (state vector **x**; Fig. 2A; Method 3.1) that evolve through recurrence (*A*), inputs (with input vector **u** and input matrix *B*), and Gaussian ‘process noise’ (**w**_*t*_ ∼ 𝒩 (0, *W* )). The observed RNN hidden states (**y**) were ‘read out’ from the latent factors according to an observation matrix (*C*) and Gaussian ‘observation noise’ (**v**_*t*_ ∼ 𝒩 (0, *V* )).

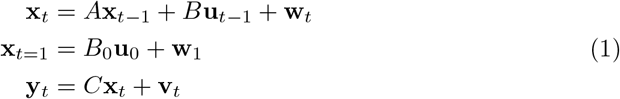

**Fig. 2:**
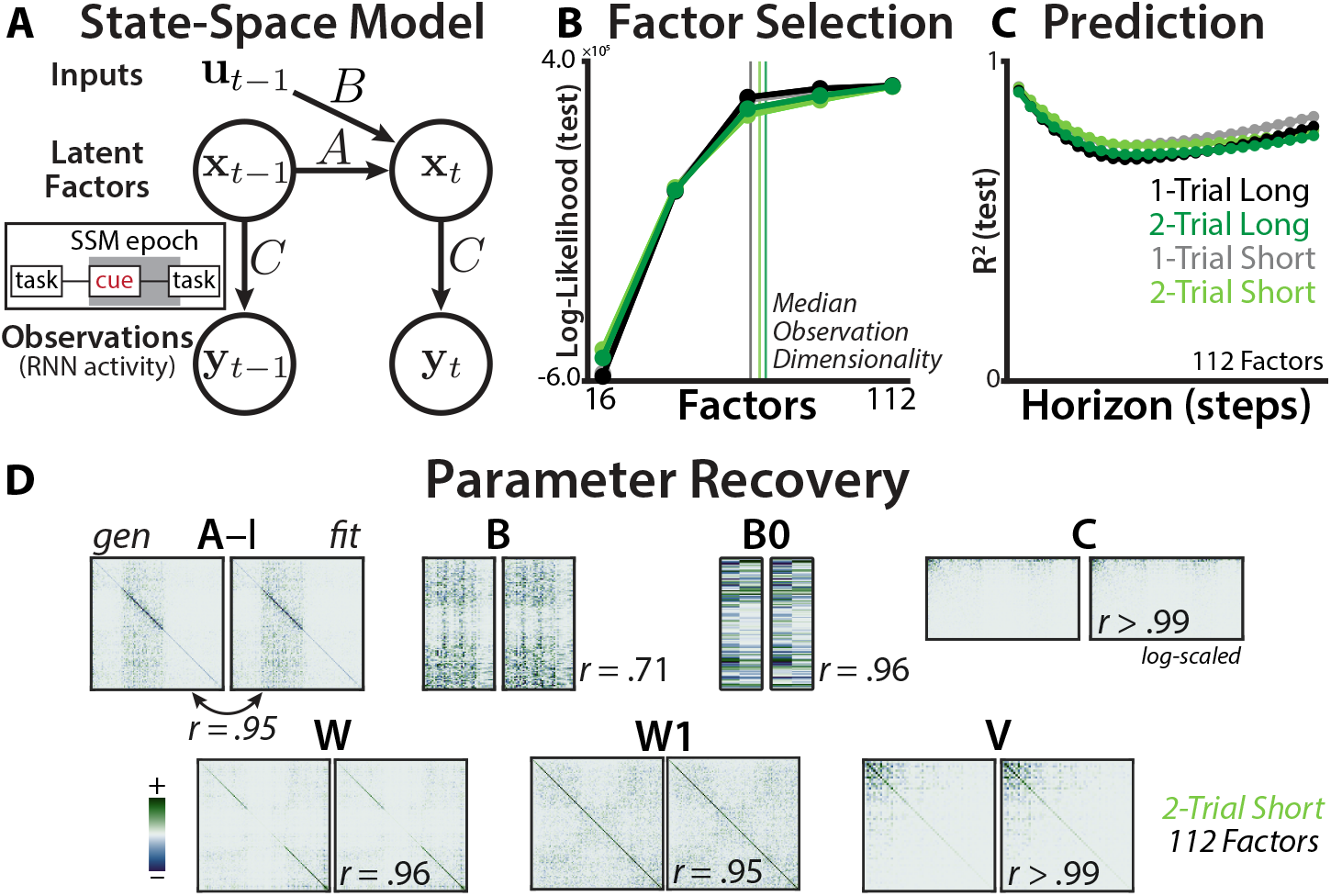
State-space analysis of RNN dynamics. **(A)** Schematic of the state-space model. Observations (**y**) are generated from latent factors (**x**), which evolve according to their intrinsic dynamics and inputs (**u**). Inset: SSM epoch included the cue period, the delay period, and the onset of the task. **(B)** SSM log-likelihood in held-out trials at different number of factors. Log-likelihood is mean-centered within each network to show relative differences. Similar fits across curricula (refer to legend in panel C). **(C)** SSM *R*^2^ in held-out trials, at different prediction horizons. Similarly high predictive accuracy across curricula, even at a horizon of 25 timesteps (more than half the epoch). **(D)** Ground truth (generating) and recovered (fit) parameters for an example RNN dataset. Parameters are linearly aligned to account for degeneracy in the latent system (Method 4.4). Colormaps are scaled separately for each parameter pair, symmetrically around zero.

We captured task-switching dynamics by including both the previous and current task identity in the SSM inputs (see Method 4.1 for the complete predictor list). To provide the model with additional flexibility, we expanded the inputs with a temporal spline basis. To capture the influence of the previous trial’s task state, the previous task identity was included in the predictors for the initial condition (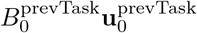; Method 4.1).

We reduced the dimensionality of the RNN state using principal component analysis (PCA), retaining PCs that accounted for 99% of the variance (between 49-76). We fit the SSM to an epoch encompassing the cue period, delay period, and the trial onset. We developed an analysis package for estimating the maximum *a posteriori* SSM parameters through a combination of subspace identification and expectation-maximization ([49–51]; Method 3).

We found that the best cross-validated fit came from ‘over-parameterized’ SSMs, which had a greater number of factors (112) than hidden units (108; Fig. 2B). This ‘lifting’ of the dimensionality allows a linear model to better approximate an underlying nonlinear system [52]. Consistent with this principle, the best-fitting model had high predictive accuracy for future timepoints on held-out trials (Fig. 2C), and had better cross-validated accuracy than autoregressive encoding models without latent states (*R*^2^ vs. null: 95% CI [.16, .30]; Method 5.2). We further validated the estimation procedure by fitting an SSM to a synthetic dataset, finding that the procedure had good recovery of the ground-truth parameters (Fig. 2D; Method 4.4), supporting the ability to differentiate effects between parameters.

### RNNs learn active reconfiguration

Having validated the predictive power of state-space modeling, we next characterized how switch-training and trial-spacing altered RNNs’ latent dynamics. Visualizing RNNs’ hidden-unit activations using ‘model-agnostic’ singular value decomposition (SVD; Fig. 3A-D), we found that all RNNs exhibited similar first-trial dynamics, but their second-trial dynamics were qualitatively different across curricula. These visualizations revealed three dynamic signatures of RNNs’ task-switching strategies, which we quantified using the fitted SSMs.

**Fig. 3:**
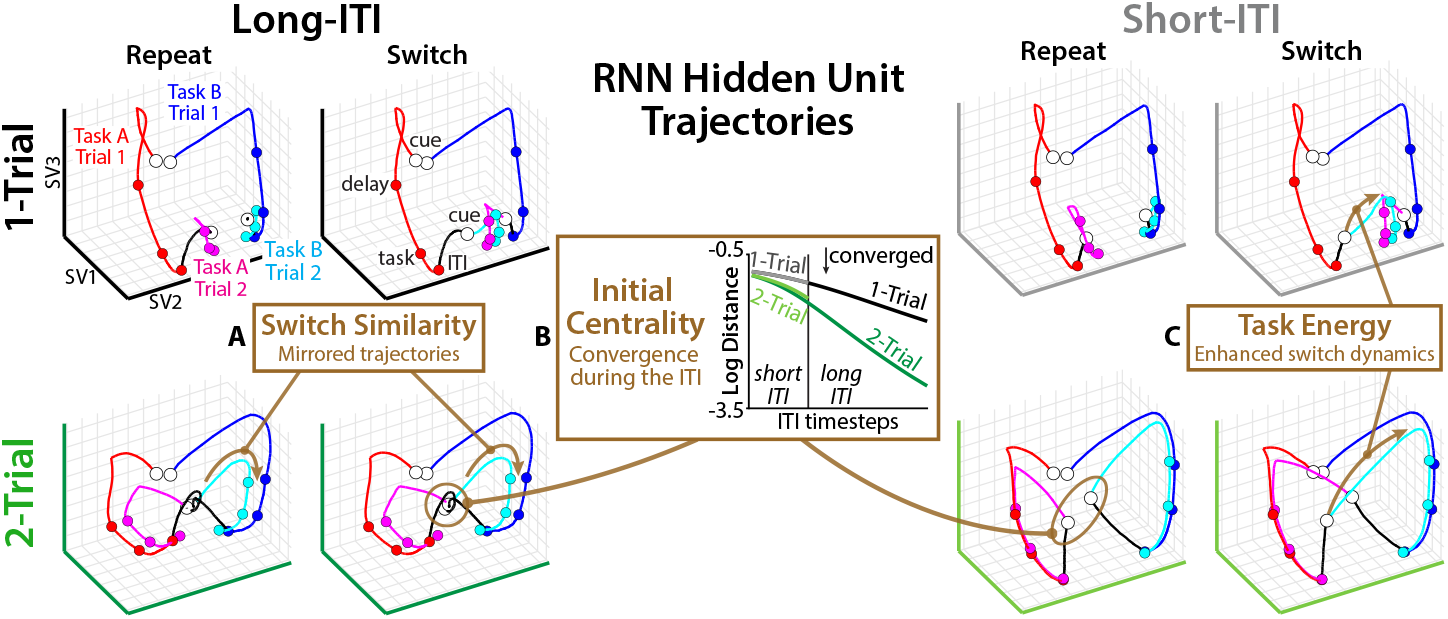
Dynamic signatures in RNN hidden unit activity. Low-dimensional embedding of example RNN hidden-unit activity for different task transitions. White dots indicate cue onset, colored lines indicate trial type, black line indicates ITI. SV: singular vector. **(A)** *Switch Similarity* : Mirrored trajectories across Switch and Repeat trials in 2-Trial RNNs, unlike 1-Trial RNNs. **(B)** *Initial Centrality* : Convergence towards a neutral state during the ITI in 2-Trial RNNs, unlike 1-Trial RNNs. Inset: Euclidean distance between RNN hidden states following each task, over the course of the ITI. **(C)** *Task Energy* : Stronger switch-dependent dynamics in 2-Trial RNNs than 1-Trial RNNs.

The first dynamic signature was the similarity of the trajectory across switch and repeat trials (Fig. 3A, ‘Switch Similarity’). On switch trials, 1-Trial RNNs appeared to directly move towards the terminal task states of the alternate task, creating opposing trajectories between switch and repeat trials. By contrast, 2-Trial RNNs’ trajectories appeared to be quite similar across switch and repeat trials. Starting from a ‘neutral’ state between the two task states, the second trial consistently reproduced the first-trial trajectories.

To quantify this first dynamic signature with our fitted SSMs, we leveraged the linearity of our SSMs to decompose the SSM into separate subsystems for each task (‘additive state decomposition’; [53]; Method 6.1):

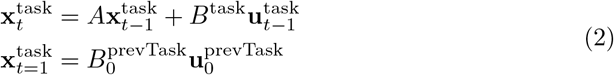

We compared the task trajectories across switch and repeat trials by computing the cosine similarity between their task states at different temporal lags ([54, 55]; Fig. 4D; Method 6.2). 1-Trial RNNs had persistent negative alignment between switch and repeat trials, consistent with opposing dynamics along a ‘task’ axis. By contrast, 2-Trial RNNs had positive alignment between switch and repeat trials, though this varied as a function of trial spacing. Whereas 2-Trial RNNs with long ITIs exhibited this positive similarity throughout the trial, those with short ITIs only showed this partway through the delay period.

**Fig. 4:**
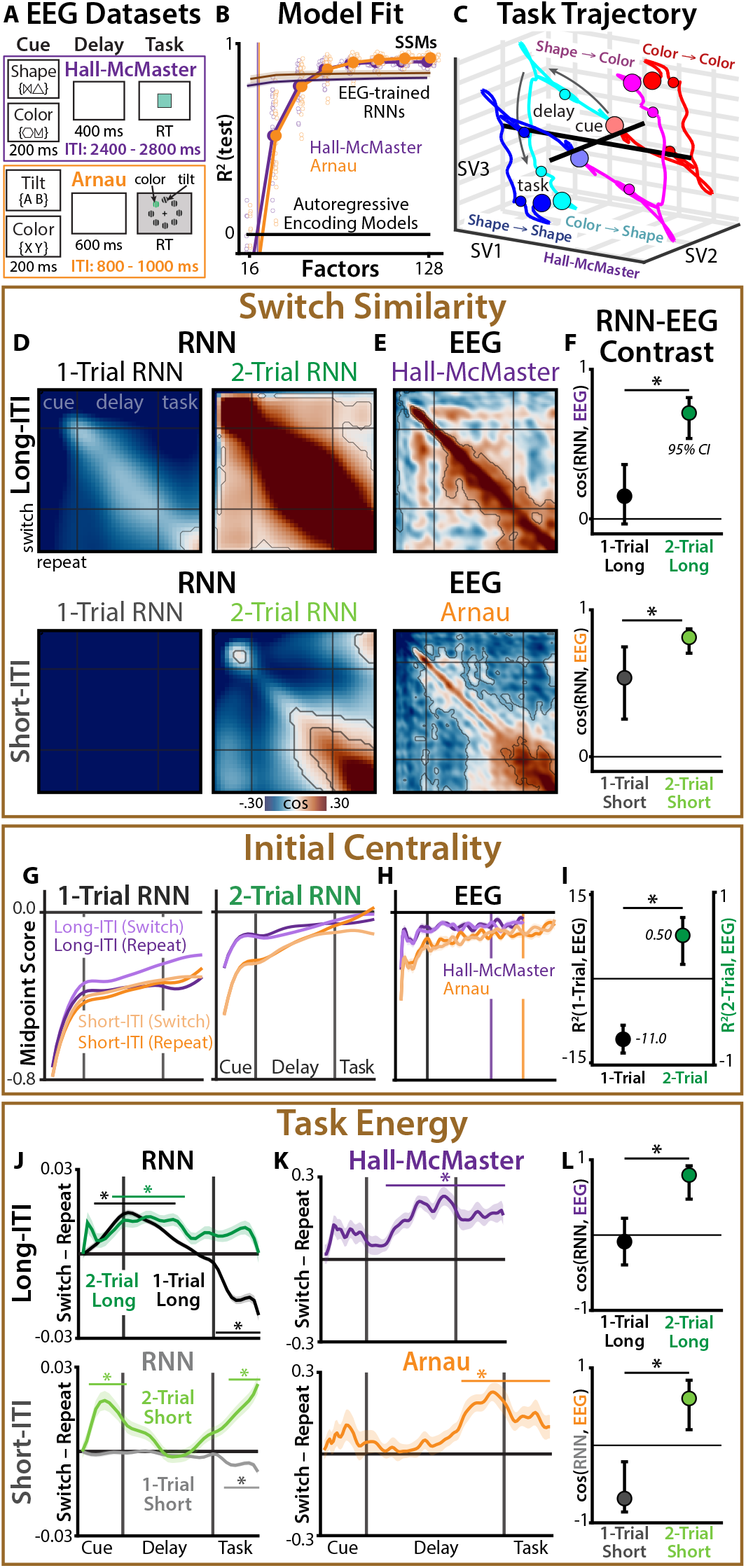
Comparison of dynamic signatures in RNNs and EEG. **(A)** Task protocol for the ‘Hall-McMaster’ (purple) and ‘Arnau’ (orange) datasets. **(B)** Cross-validated *R*^2^ at different number of latent factors. Zero indicates fit of autoregressive encoding model. Darkly colored lines indicate EEGtrained RRN models. Vertical lines indicate median observation dimensionality. **(C)** Low-dimensional embedding of SSM task trajectories, split by task transitions (SV: singular vector). Task is contrast-coded, so trajectories are symmetrical within switch condition. Shown for Hall-McMaster, see Fig. S3 for Arnau. **(D-E)** *Switch Similarity* : Cross-lagged similarity between SSM task states on switch vs. repeat trial, for (D) RNNs and (E) EEG. Cardinal lines indicate boundaries between cue, delay, and task events. Contours indicate *p <* .05, corrected for multiple comparisons with ‘threshold-free cluster enhancement’ (TFCE; Method 6.5). **(F)** Bootstrapped cosine similarity between EEG and 1-Trial vs. 2-Trial RNNs. **(G-H)** *Initial Centrality* : relative distance between SSM initial conditions and each task states, for (G) RNNs and (H) EEG. Zero indicates initial conditions are at the midpoint. **(I)** Bootstrapped *R*^2^ between EEG and 1-Trial vs. 2-Trial RNNs. Note that y-axis have different scales for better visualization. *R*^2^ metric can have negative values (Method 5.2). **(J-K)** *Task Energy* : difference in task energy between switch and repeat trials, for (J) RNNs and (K) EEG. Asterisks indicate *p <* .05 (TFCE-corrected). **(L)** Bootstrapped cosine similarity between EEG and 1-Trial vs. 2-Trial RNNs.

The second dynamic signature was the convergence of task states towards a neutral state during the ITI. Starting from distinct previous-task states, the distance between states during the ITI decreased as they moved into a common state (Fig. 3B ‘Initial Centrality’). This convergence was faster for 2-Trial RNNs than for 1-Trial RNNs, despite having a similar initial distance.

Starting the second trial at a neutral state may explain why 2-Trial RNNs had higher switch similarity than 1-Trial RNNs: starting near the midpoint between task states produced symmetrical trajectories. It also may explain why ITI length influenced switch similarity in 2-Trial RNNs: shorter ITIs left less time to reach the neutral state.

To understand *where* RNNs had converged by the end of the ITI, we used SSMs to quantify whether the second trial’s initial conditions were near the midpoint between task trajectories. We measured the relative Euclidean distance of each task state to the initial conditions (Method 6.3):

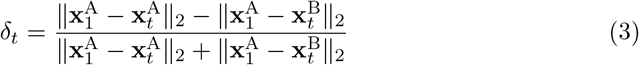

In 1-Trial RNNs, the initial state was far from the task midpoint, without strong differentiation across ITI conditions (Fig. 4G). In 2-Trial RNNs, the initial state was near the midpoint of the task state, and RNNs with longer ITIs achieved a more central position (especially for cue and delay states). This confirms that RNNs strategically transitioned into a neutral state during the ITI, one that was well-positioned for the upcoming trial phases.

The third dynamic signature revealed post-cue differences in task control. Following the task cue, 2-Trial RNNs appeared to have much more robust dynamics than 1-Trial RNNs, especially when switching between tasks (Fig. 3C, ‘Task Energy’). To quantify the extent to which RNN trajectories were driven by cue-dependent control, we used ‘Lyapunov analyses,’ a standard tool from control theory for quantifying the influence of control inputs [42]. Previous work has used these methods for asymptotic ‘controllability’ analyses, modeling how control inputs could drive a neural system in principle [43, 56, 57]. Here, we quantified the realized ‘task energy’ using a recursive Lyapunov equation [58] (Method 6.4):

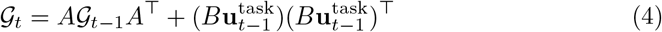

Comparing the average task energy between switch and repeat trials revealed a strong influence of switch-training (Fig. 4J). 1-Trial RNNs exhibited weak or absent differences between switch and repeat conditions, especially at short ITIs. By contrast, 2-Trial RNNs in both ITI conditions showed stronger task energy on switch trials than repeat trials. Through their training on task switching, 2-Trial RNNs learned active control strategies that were responsive to ongoing task demands.

These findings suggest an important augmentation of the standard view of how neural control processes flexibly switch between tasks. Rather than switching directly from one task state to another at the time of the cue, these results suggest that switch-trained RNNs move towards an intermediate, ‘neutral’ state before a task cue, a state that is roughly equidistant between the upcoming task states. Then, when the task cue appears, RNNs gate in this new information to drive the system toward the corresponding task state. Below, we test whether similar latent dynamics are observed in EEG recordings from the human brain during task switching.

### Analogous task control in artificial neural networks and human brains

We re-examined two previously reported human EEG datasets, collected during complementary task switching experiments. The first was from an experiment by Hall-McMaster et al. [59], in which 30 participants performed a cued task-switching paradigm while 61-channel EEG was recorded (Fig. 4A; Method 2.1). On each trial, participants were cued whether to perform a ‘shape’ or ‘color’ task. Following a delay period, participants saw a colored shape and responded with a button press based on the cued dimension. The second dataset was from Arnau et al. [60], in which 26 participants performed a similar task-switching paradigm while 128-channel EEG was recorded, here switching between color and orientation targets in a visual search task (Fig. 4A; Method 2.2).

Critically, these two datasets had substantially different ITIs, with much longer ITIs in ‘Hall-McMaster’ (2400 - 2800 ms) than ‘Arnau’ (800 - 1000 ms). We used these datasets to test the extent to which the human brain exhibits similar task dynamics to those that we observed in RNNs, including movement into a neutral task state over the ITI.

We fit SSMs to each EEG dataset, using the same fitting procedure that we used for RNNs. We again found that the best-fitting models had more latent factors than EEG components (Hall-McMaster: 112 factors, 14-27 PCs; Arnau: 128 factors, 15-35 PCs; 112 factors used throughout; Fig. 4B). As with the RNNs, over-parameterized SSMs made accurate long-range predictions in held-out data (Fig. S2) and had good parameter recovery (Fig. S4). SSMs provided better predictions of EEG activity than both simpler autoregressive encoding models and nonlinear RNNs trained to predict EEG (Fig. 4B; Method 5).

Visualizing the inferred EEG task trajectories (Fig. 4C; Fig. S3), we found that switch and repeat trials appeared to have a similar timecourse, as in switch-trained RNNs. EEG task states also appeared to support good performance: when EEG recordings were better aligned with the inferred task state, participants tended to have faster reaction times (fixed effects from linear mixed model, Hall-McMaster: *d* = 0.59, *p* = .0073; Arnau: *d* = 0.75, *p* = .012; Method 8.2).

By modeling RNN and EEG datasets within the same SSM framework, we could directly test dynamic signatures of active configuration across theory and data. Across all three signatures, we found that 2-Trial RNNs were a better match to human EEG than 1-Trial RNNs, consistent with reconfiguration. We also found that differences between RNN datasets with long vs. short ITIs were recapitulated by differences between EEG datasets with long vs. short ITIs, consistent with an influence of the previous task state.

We first compared EEG and RNNs datasets on their patterns of cross-lagged similarity between switch and repeat trials (‘Switch Similarity’; Fig. 4D). EEG exhibited similar task encoding across switch and repeat trials, which was more consistent with the pattern observed in 2-Trial RNNs than 1-Trial RNNs (contrasting 2-Trial vs. 1-Trial cosine similarity with EEG: Hall-McMaster: 95% CI [0.18, 0.83], Arnau: 95% CI [0.076, 0.46]; Fig. 4E-F; Method 7).

In 2-trial RNNs, we had found that positive switch similarity arose from convergence towards a neutral task state during the ITI (‘Initial Centrality’; Fig. 4G). This pattern was closely reproduced in the two EEG datasets (contrasting 2-Trial vs. 1-Trial *R*^2^ with EEG: 95% CI [8.3, 13]; Fig. 4H-I), including more central initial conditions in the longer-ITI dataset (two-sample t-test on midpoint score, averaged over cue period: *d* = 1.16, *p* = 6.54 × 10^-5^; averaged over delay period: *d* = 0.75, *p* = .0068).

Finally, we had found that 2-Trial RNNs exhibited greater cued task energy on switch trials than on repeat trials (‘Task Energy’; Fig. 4J). Here too, EEG datasets reproduced 2-Trial RNNs’ pattern of switch-elevated task control (contrasting 2-Trial vs. 1-Trial cosine similarity with EEG: Hall-McMaster: 95% CI [0.87, 1.2]; Arnau: 95% CI [1.3, 1.6]; Fig. 4K-L). Together, these findings demonstrate a striking correspondence between task-optimized RNNs and human neural dynamics, revealing a common set of strategies for flexible information processing.

## Discussion

While psychologists have debated for decades over the latent processes underlying cognitive flexibility [2, 3], a quantitative account of the underlying neural mechanisms has remained elusive. To provide such an account, we trained neural networks on curricula that manipulated the primary psychological factors that are theorized to influence task-switching performance: active preparation through cognitive control and proactive interference from prior tasks. To quantify how these factors affected RNN dynamics, we fit state-space models to the RNN hidden unit activity. This state-space modeling approach allowed us to precisely compare control dynamics in RNNs and human brain recordings using a common statistical model.

These analyses revealed that RNNs trained to switch tasks deploy a control strategy in which they transition into a neutral task state during the ITI, like a tennis player recovering to the center of the court after a shot. This represents a novel augmentation to existing task-switching theories, reconciling the effects of previous task states with active reconfiguration. It takes time to reconfigure into the neutral state, which makes switch and repeat dynamics more symmetric at longer inter-trial intervals. Critically, this strategy depended on RNNs being trained to switch between tasks, consistent with it reflecting a learned strategy.

We tested the extent to which this RNN model could explain human brain activity during task switching. We deployed the same inferential model and control theoretic analyses on two EEG datasets, which we then compared to the trained RNNs. These analyses revealed that human brain activity exhibits strikingly similar dynamics to those observed in RNNs that learn control strategies for switching between tasks. In particular, the human brain appears to replicate the RNN strategy of moving into a neutral state between trials, highlighting a potential benefit to ‘mentally resetting’ between difficult tasks.

These findings provide a valuable framework for understanding the neuro-computational mechanisms underlying classic psychological theories of task-switching. We find strong evidence that humans actively prepare for an upcoming task [12]. At the same time, we also find strong evidence that task switching depends on prior task states [14]. The influence of both factors can be explained within a unifying control framework that emerged from the optimization of task performance in gated neural networks.

These theoretical insights were enabled by the capabilities of linear-Gaussian state-space models to approximate nonlinear systems like RNNs and human brains. In the over-parameterized regime (more latent than observed dimensions), SSMs exhibited a powerful combination of predictive power and interpretability. While this state-space approach has considerable promise for human neuroscience, some caveats remain to be addressed. First, the full expressivity of our dynamics came from modeling time-varying inputs using a descriptive model (spline basis) rather than a process model. Future research should model the task inputs (e.g., using optimal control [43, 56, 61, 62]). A second caveat is that switch-trained RNNs showed weaker switch costs than expected from human behavior (Fig. S1), despite similar preparatory neural dynamics. This is consistent with theories that posit additional contributions to switch costs beyond preparation (e.g., associative learning, [11, 17, 63]), and future research should extend these findings to better accommodate behavioral switch costs [23, 26]. Caveats aside, the approaches utilized here, marrying neural network modeling, latent inference, and control theory, have the potential to help scientists better understand the dynamics of distributed neural systems that enable flexible cognition, both *in vivo* and *in silico*.

## Acknowledgments

We are extremely grateful to Sam Hall-McMaster and Stefan Arnau for making their datasets public and providing additional support. Thanks to Christopher Langdon for sharing PyTorch code for context-dependent decision-making tasks, Caroline Jahn for feedback on an early draft of the manuscript, Jonathan Pillow for helpful advice on inferential methods, and the Daw and Cohen labs for their feedback throughout. This paper is dedicated to Mark Stokes.

## Funding

HR was supported by the C.V. Starr Postdoctoral Fellowship. AJ was supported by the Google PhD Fellowship. JDC was supported by a Vannevar Bush Faculty Fellowship administered by the Office of Naval Research.

## Author contributions

Conceptualization: HR, NDD, JDC

Data curation: HR

Formal analysis: HR, AJ

Funding acquisition: HR, NDD, JDC

Methodology: HR, AJ

Resources: NDD, JDC

Software: HR

Supervision: NDD, JDC

Writing – original draft: HR

Writing – review & editing: HR, AJ, NDD, JDC

## Competing interests

There are no competing interests to declare.

## Data and materials availability

Paper analysis code: www.github.com/harrisonritz/SSM-Paper-2025

State-space analysis package: www.doi.org/10.5281/zenodo.14511205

‘Hall-McMaster’ dataset: www.osf.io/kuzye/

‘Arnau’ dataset: www.osf.io/ndgst/

## Methods

### 1 Task-Optimized Recurrent Neural Networks

#### 1.1 RNN Architecture

We trained RNNs with gated recurrent units (GRUs; 108 hidden units; [27]) to perform task-switching. These networks used a standard single-layer RNN architecture: linearly encoded inputs (*x*_*t*_), a recurrent hidden state (*h*_*t*_) with a hyperbolic tangent activation function, and linearly decoded outputs (*y*_*t*_) with a sigmoid link function.

The GRU component of this RNN learned two ‘gates’, each of which encodes the hidden state and inputs, and outputs a sigmoid-constrained gating value that is element-wise multiplied with the hidden state. The ‘reset gate’ (*r*_*t*_) is applied to the hidden state before the activation function, producing the ‘new state’ (*n*_*t*_). The ‘update gate’ (*z*_*t*_) is applied after the activation function, mixing the new state and the previous hidden state. This was implemented through the standard GRU class in PyTorch [64]:

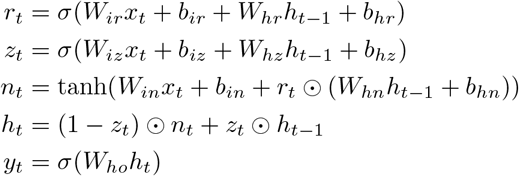

Note that for consistency with PyTorch, we use *h*_*t*_ to refer to the latent state (SSM: *x*_*t*_), and *x*_*t*_ to refer to the inputs (SSM: *u*_*t*_). We trained the GRUs using backpropagation through time, using the AdamW learning rule (learning rate = .01, weight decay = .01).

#### 1.2 Curricula

We trained four groups of GRUs on different curricula (512 RNNs per curriculum, with matched random seeds), crossing two levels of ‘Switching-Training’ against two levels of ‘Trial-Spacing’.

##### 1.2.1 Switch-Training

‘1-Trial’ RNNs were trained on sequences containing a single trial (9600 sequences *×* 500 epochs; see task description below). ‘2-Trial’ RNNs were trained on sequences of one trial for 90% of their training (450 epochs), and then trained on sequences of two trials for the final 10% of training (50 epochs). Each epoch had an equal number of target transitions, distractor transitions, and task transitions (64 conditions × 150 repetitions). The rationale for this design was to teach both groups how to perform the task, but give 2-Trial networks enough experience that they would learn a compensatory strategy for switching. By 450 epochs, RNNs were over-trained on the task, with increasing test loss (Fig. 1E).

##### 1.2.2 Trial-Spacing

‘Short ITI’ RNNs were trained on sequences in which the inter-trial interval between trials was short (20 timesteps; see task description below), whereas ‘Long ITI’ RNNs were trained on sequences with longer ITIs (60 timesteps). This ratio of ITIs was chosen to match the EEG experiments (see below). For 1-Trial networks, this ITI manipulation only affected their test trials.

#### 1.3 Task

RNNs performed an extension of the ‘context-dependent decision making’ task (CDM; [28, 29, 65, 66]. On one-trial sequences, RNNs performed a classic CDM task. In each sequence, RNNs first received a task cue (one-hot coded; 10 timesteps), followed by a delay period (20 timesteps). During the task period, they got two pairs of stimulus inputs (40 timesteps). They had to judge which input in the task-relevant pair (i.e., as indicated by the task cue) had a higher amplitude. On two-trial sequences, RNNs performed two consecutive trials (i.e., without resetting the state or gradient). They performed the first trial, had an ITI with no inputs (20 or 60 timesteps, see above), and then performed the second trial.

RNNs were trained to optimize a logistic loss function on the correct (*y*_*i*_ = 1) and incorrect (*y*_*i*_ = 0) response options ( ℒ = − ∑_*i*_(*y*_*i*_ log(*ŷ*_*i*_) + (1 − *y*_*i*_) log(1 − *ŷ*_*i*_))), with the loss starting 10 timesteps after the onset of the stimuli to encourage evidence integration. We added normally distributed noise to the inputs in each sequence (cue SNR: 0 dB; trial SNR: -16.5 dB), which was resampled every epoch. We trained at relatively low SNRs to encourage the networks to use task control and to discourage overfitting. The test loss was evaluated on the 64 conditions, presented without noise. To provide a stronger test of generalization, we also evaluated the trained RNNs on novel three-trial sequences (Fig. 1D).

#### 1.4 Gate Ablations

To assess the role of the reset and update gates in GRU’s performance, we trained a series of ablated models. These ablated models consisted of (1) fixing the reset gate to be open (full use of previous hidden states), (2) fixing the update gate to be open (full use of new state), or (3) a vanilla RNN (no gating). GRUs were trained with tanh activation function and learning rate = .01, whereas the RNN was trained with a ReLU activation function and learning rate = .001 (which improved its performance). Free parameters were matched across networks (108 hidden units for the full model, 134 units for ablated GRUs, 190 units for RNN). To assess the influence of trained these models on 500 epochs of one-trial sequences, and then 500 epochs of two-trial sequences.

#### 1.5 Hidden Unit Visualization

We used singular value decomposition (SVD) to visualize hidden unit activation during two-trial sequences. Using trained networks, we first simulated a set of noisy trials (n=2048) for an example network from each curriculum (same random seed). Next, we concatenated the hidden unit activations across sequences and networks into a 2D matrix ((timesteps *×*sequences) *×* (hidden units *×* curricula)). Performing SVD on this matrix provided embeddings that captured the shared and distinct components of each network, which was qualitatively similar to when we used a single embedding for all RNNs. We then averaged the sequences within each network and task transitions, and projected these average sequences onto the right singular vectors corresponding to each network.

### 2 EEG Datasets

#### 2.1 Hall-McMaster Dataset

##### 2.1.1 Sample

The first EEG dataset was originally published by Hall-McMaster and colleagues [59], and was made open-access at www.osf.io/kuzye/. Thirty participants (18-35 years old; 19 females) with normal or corrected-to-normal vision and no history of neurological or psychological disorders underwent an EEG recording session during task performance. This study was approved by the Central University Research Ethics Committee at the University of Oxford, with all participants providing informed consent. See [59] for the full description.

##### 2.1.2 Task

Participants performed a cued task-switching experiment during the EEG recording. A central goal of the original study was to investigate the role of incentives on task encoding, a manipulation which was not the focus of the current experiment.

Participants saw a cue for high vs. low rewards for correct performance (800 ms, 400 ms delay), a cue for the shape vs. color task (200 ms, 400 ms delay), a task stimulus consisting of a square or circle colored in yellow or blue (min(RT, 1400 ms)), reward feedback (200 ms), and an inter-trial interval (1000-1400 ms). Participants’ task was to respond to the trial stimulus according to the cued rule: if the task was ‘shape’, respond with a key press based on the shape; if the task was ‘color’, respond with a key press based on the color. Participants practiced the task until they reached a 70% accuracy criterion, and then performed 10 blocks of 65 trials in the main task. Conditions were balanced within each block.

##### 2.1.3 EEG Preprocessing

For our analyses, we included trials 1) that were not the first trial of a block, 2) where the current and the previous trial were accurate, 3) where the current RT was longer than 200 ms, and 4) that were not identified in the original experiment as containing artifacts during preprocessing. We epoched the data between the onset of the task cue and 200 ms into the task. To avoid removing previous task representations, we did not apply any baseline correction.

Working with the preprocessed data from the original experiment, we low-pass filtered the EEG at 30 Hz (resulting in a 0.01 Hz - 30 Hz bandpass filter) and then resampled the data to 125 Hz using Fieldtrip. We split up the experimental blocks into training and test sets, with Mean(SD) = 416(63) training trials and 47(9) testing trials. EEG data were recorded with 61 Ag/AgCl sintered electrodes (EasyCap; 10-10 layout (Fieldtrip template: elec1010)), a NeuroScan SynAmps RT amplifier, and using Curry 7 acquisition software.

#### 2.2 Arnau Dataset

##### 2.2.1 Sample

The second EEG dataset was originally published by Arnau and colleagues [60], and was made open-access at www.osf.io/ndgst/. This dataset consists of twenty-six participants (23 female, mean(SD) age 21.65(2.15) years), with normal or corrected-to-normal vision and no history of neurological or psychological disorders, and verified normal color vision. This study was approved by the ethics committee of the Leibniz Research Centre for Working Environment and Human Factors, Dortmund, with all participants providing informed consent. See [60] for the full description.

##### 2.2.2 Task

Participants performed a cued task-switching experiment during the EEG recording. On each trial, participants saw a cue for the tilt vs. color task (200 ms, 600 ms delay). During the task phase, participants saw an array of Gabor patches and had to report either the tilt or the color of the singleton, depending on the cued task. Participants had up to 1200 ms to respond, and then there was an inter-trial interval (800-1000 ms). As in Hall-McMaster, participants practiced the task before the main experiment, which consisted of eight experimental blocks with 256 trials each. This experiment also investigated the role of incentives on task switching using a mini-block-level incentive manipulation. This similarity is coincidental, due to motivation being an active area of focus in recent high-quality experiments on cognitive control.

##### 2.2.3 EEG Preprocessing

EEG data were recorded using 128 passive Ag/AgCl electrodes (Easycap GmbH, Herrsching, Germany) with a NeurOne Tesla AC-amplifier (Bittium Biosignals Ltd., Kuopio, Finland). We used the same performance-based inclusion criteria as in Hall-McMaster. EEG data were preprocessed using EEGLab scripts modified from the original experiment. We high-pass filtered at 0.1 Hz and low-pass filtered at 30 Hz using FIR filters, and then resampled the data at 125 Hz. To avoid removing previous task representations, we did not apply any baseline correction. We used more lenient inclusion criteria for the ICA auto-rejection procedure than the original experiment, as we found that some participants had most of their components removed. After preprocessing, we split up the experimental blocks into training and test sets, with Mean(SD) = 676(147) training trials and 142(31) testing trials.

### 3 Linear-Gaussian State-Space Model

#### 3.1 Generative Model

We developed a toolkit called StateSpaceAnalysis for fitting linear-Gaussian state-space models (SSMs) to neuroimaging data, inspired by the SSM and dynamax packages in python [67]. Our goal was to test how effectively SSMs could capture rich neuroimaging data, balancing the predictive power of machine learning approaches with the interpretability offered by linear-Gaussian system identification. To provide the speed and accuracy needed for fitting high-dimensional latent variable models, we developed this package using the powerful open-source coding language julia, available at www.github.com/harrisonritz/StateSpaceAnalysis.jl [51].

The generative model for the analysis is a partially observable autoregressive process: a linear dynamical system from which we can only make noisy measurements (or, equivalently, that only produces noisy emissions of its underlying state). This autoregressive process consists of a vector of latent factors (**x**), and a discrete-time difference equation that describes how they evolve:

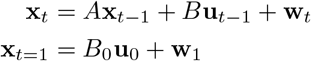

Latent factors evolve according to their recurrent dynamics (*A***x**), the influence of known inputs (*B***u**), and process noise (**w**_*t*_ *∼* 𝒩 (0, *W* )). The initial conditions depend on trial conditions (*B*_0_**u**_0_) and initial uncertainty (**w**_1_ *∼* 𝒩 (0, *W*_1_)).

At each timestep, we get a noisy observation of these latent factors:

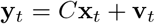

Observations are projections of the latent factors (*C***x**) corrupted by observation noise (**v**_*t*_ *∼* 𝒩 (0, *V* )). Note that only process noise is carried forward in time through the recurrent dynamics, unlike observation noise.

#### 3.2 Expectation-Maximization

A major benefit of linear-Gaussian SSMs is that they allow closed-form maximum *a posteriori* inference of the underlying latent state, and closed-form marginal likelihood of the observed data, *P* (**y**_1:*T*_ | Θ) where Θ represents the model parameters. This is achieved using standard inferential tools: Kalman filtering (inferring **x**_*t*_ from past observations) and Rauch–Tung–Striebel (RTS) smoothing (inferring **x**_*t*_ using the entire trajectory) for latent estimation (E-Step), and linear regression for parameter inference (M-Step).

While the marginal likelihood from Kalman filtering would allow us to directly fit the parameters through techniques like maximum likelihood estimation, in practice it is more efficient to use expectation-maximization (EM; [49, 50, 68]). EM maximizes the expected lower bound of the marginal posterior (often called the evidence lower bound or ELBO) by alternating between two steps. In the E-step, we estimate the latent state using the RTS smoother. In the M-step, we find point-estimates of the parameters that maximize the ELBO, given the estimated latent states. With priors on the parameters, this estimates the mode of their posterior density. The Linear-Gaussian assumptions of this model considerably simplify this procedure by allowing us to work with the sufficient statistics of the estimated latent state (i.e., their mean and covariance).

##### 3.2.1 Expected Log-Posterior

Our M-Step maximized the expected log-posterior (i.e., the ELBO) over *N* epochs and *T* timesteps:

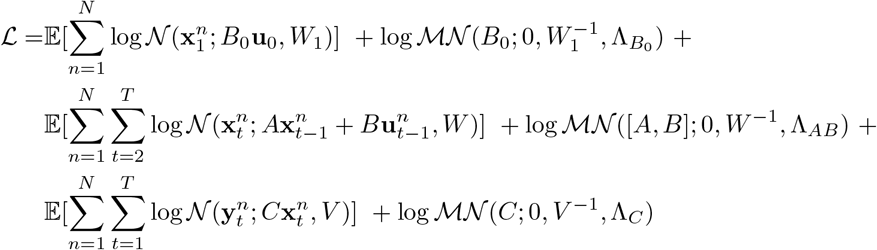

We used weakly informative priors for [*A, B, B*_0_, *C*], with prior means of zero and prior precisions set to scaled identity matrices (Λ = 10^−6^*I*). We used uninformative priors for [*W, W*_1_, *V* ], which are excluded from above.

##### 3.2.2 E-Step

Using the filter-smoother algorithm [69], we estimated the posterior mean (**m**_*t*_) and covariance (Σ_*t*_) of the latent state across *N* epochs and *T* timesteps, generating a set of sufficient statistics:

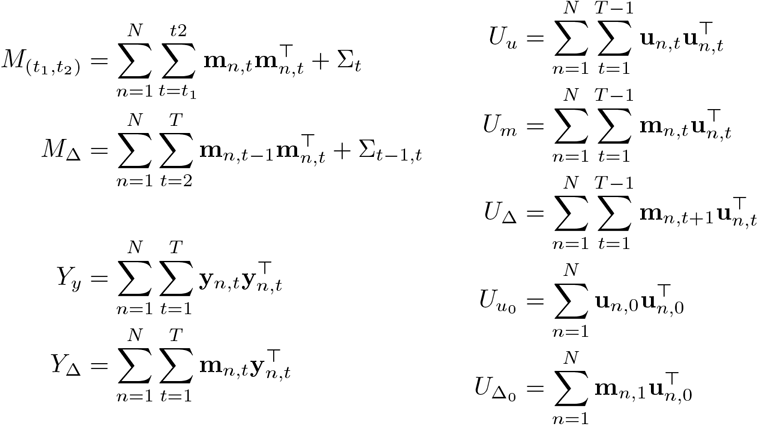

Note that we can use the same smoothed covariance estimates (Σ) for all epochs, substantially speeding up the computation.

##### 3.2.3 M-Step

We then use the computed sufficient statistics to update the parameters using a maximization procedure similar to ordinary least squares, with the inclusion of ridge priors on all of the dynamics matrices (Λ).

For the dynamics terms:

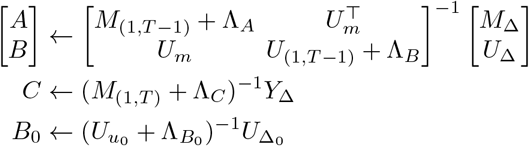

For the covariance terms:

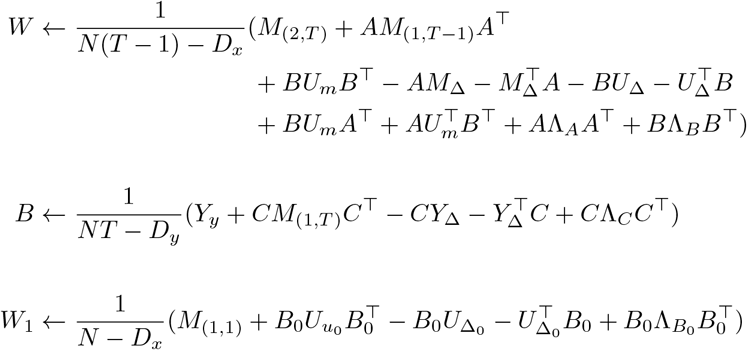

With *D*_*x*_ and *D*_*y*_ indicating the numbers of factors and observation dimensions, respectively. Note that the estimator of *W*_1_ is unusual, as the initial conditions are not usually parameterized with inputs, however, this is equivalent to the standard formula from [49] when the only input is an intercept. Alternating between these E-steps and M-steps monotonically improves the ELBO towards a local optimum, with a strong dependence on the initialization (see ‘Subspace Identification’ below).

#### 3.3 Subspace Identification (SSID)

We found that good initialization of the parameters was critical to the effectiveness and stability of the model-fitting procedure. We initialized the parameters using a procedure called subspace identification (SSID; [70, 71]). This procedure constructs a large delay embedding matrix which concatenates lagged future and past copies of the observations (**y**) and inputs (**u**). The ‘horizon’ of these lags was set to match the number of factors in the largest tested model (see below). We used the canonical variate analysis procedure [72], which uses SVD to find low-dimensional mappings between lagged inputs and outputs. We can recover the *A* and *C* matrices by processing different blocks of this SVD, and then estimate the rest of the parameters using infinite-horizon Kalman filtering. We performed CVA using a modified version of the ControlSytemIdentification.JL Julia package (Method 9).

### 4 Fitting procedure

#### 4.1 SSM Preprocessing: RNN

The fitting procedure was closely matched between networks and EEG. We separately preprocessed RNN training and test trials (training: 1536 trials; testing: 512) by performing PCA on the hidden unit trajectories (keeping components accounting for 99% of the variance; estimated on training data only) and constructing a temporal basis for the inputs (10 spline bases over the epoch). Inputs included a (time-varying) bias, the cued task, the previous trial’s task, switch vs. repeat. Inputs to initial conditions were an intercept and the previous task. Each input was z-scored across trials to standardize and reduce collinearity. We analyzed the preparation for the second trial in each sequence: cue period (10 timesteps), delay period (20 timesteps), and the initial trial period (10 timesteps).

We fit across five levels of latent dimensionality (16, 40, 64, 88, 112), initializing the parameters with SSID at a horizon of 112. We fit the SSM to all 512 1-Trial and 2-Trial RNNs.

#### 4.2 SSM Preprocessing: EEG

We followed similar procedures for the Hall-McMaster and Arnau datasets as we did in the RNNs. We split the data into training and testing folds, preprocessing the inputs and observations of each separately. We epoched the recordings around the preparatory period, including the cue (200 ms), the delay period (Hall-McMaster: 400 ms; Arnau: 600 ms), and the first 200 ms of the trial (which we ensured was before any RTs through our trial rejection).

The SSM inputs included a (time-varying) bias, the current task, the previous task, the cue identity (with separate regressors contrasting the cue symbols within each task), cue repetitions vs. alternations for repeat trials, task switch vs. task repeat, the upcoming trial’s reaction time, and the previous trial’s reaction time. For the initial conditions, we included an intercept, the previous task, the previous reaction time, and the upcoming reaction time. We expanded the predictors with a cubic B-spline basis set. Spline knots were placed to optimize coverage at every 5 timesteps, tiling the entire epoch (Hall-McMaster: 20 bases per predictor, Arnau: 25 bases per predictor). Each input was z-scored across trials to standardize and reduce collinearity.

We preprocessed the EEG electrodes using PCA, projecting the voltage timeseries into PCs accounting for 99% of the variance. We used PCA because the observations were degenerate due to spatial proximity, independent components analysis, and bad-channel interpolation. PCA also reduced the computational demands of these analyses by reducing the observation dimensionality and making the observation covariance closer to diagonal. We estimated the PCs using only the training data, and then projected the test data onto these PCs.

As described above, the EM procedure cycled between estimating the latent states and updating the parameters. For EEG datasets, we fit the model across eight different levels of latent dimensionality: (16, 32, 48, 64, 80, 96, 112, 128). We set the maximum number of iterations to 20,000, measuring the test-set log-likelihood every 100 iterations. We terminated the fitting procedure if either the total data likelihood in the training set stopped decreasing, or if the test log-likelihood stopped decreasing.

#### 4.3 SSID Procedure

To provide the best initialization across the latent dimension hyper-parameter, we estimated SSID for a horizon and latent dimensionality that matched the largest tested model (128 latent factors). We then truncated these systems for smaller models (as latent states from CVA are ordered by their singular values). We found that SSID was enhanced by reshaping the trial-wise data into a long timeseries (e.g., allowing for larger numbers of factors and low frequencies). During SSID, we also only included inputs in the initial timesteps of each epoch, reducing the collinearity between lagged inputs. Note that we removed the test set before SSID (which was on a separate experimental blocks to minimize temporal proximity). The EM procedure, unlike SSID, was not fit on concatenated epochs and used all of the inputs. In practice, we found that this SSID procedure provided a good initial guess of the parameters while avoiding requiring multiple initializations, and was especially important when there were more factors than observation dimensions.

#### 4.4 Parameter Recovery

We validated that the combination of model, data, and fitting procedure could produce identifiable parameter estimates under a known generative model. First, we used the SSM generative model to create a synthetic dataset based on a participant’s parameters and inputs. We then estimated the parameters of this synthetic dataset using EM. Next, we aligned the estimated and recovered parameters, as the estimates from are only identifiable up to an invertible transformation (*T* = Diagonal(*W* ); *A*^*′*^ = *T* ^−1^*AT* ; *B*^*′*^ = *T* ^−1^*B*; *C*^*′*^ = *CT* ; see [42] § 2.5). Intuitively, we could shuffle the ordering of the latent factors without changing the likelihood. Finally, we correlated the generating and recovered parameters (correlating the lower triangle of the Cholesky factors for covariance matrices).

Our recovery was overall very high, but we had somewhat poorer recovery of input matrices. We found that this strongly depended on the norm of the input matrix column that we were trying to recover: some predictors were weakly encoded, and these were difficult to recover. It also likely reflects the correlation between proximal temporal bases.

### 5 Model Comparison

#### 5.1 Bayesian Model Selection

We selected the number of latent factors for subsequent analyses using a standard Bayesian model selection procedure [73]. This analysis estimates the expected probability of each model in the population. From these posterior mixture weights, we computed a ‘protected exceedance probability,’ the probability that a model is the most popular within the tested set of models, relative to chance. While the best-fitting model in the Arnau dataset had 128 factors, we used the second-best model (112 factors) to allow for fair comparisons with Hall-McMaster and RNN models.

#### 5.2 Sensor-Level Null Models

To compare to a set of simpler null hypotheses, we fit a set of four models directly to the electrode PCs (i.e., without a latent embedding beyond the PCA). First was an intercept-only model (i.e., the standard *R*^2^). Second was an encoding analysis using the full temporal basis set (a generalized additive model similar to EEG methods like the unfold toolbox, [74]). Third was a vector autoregressive (VAR(1)) model, estimating the multivariate relationship between sensor PC responses on adjacent timesteps. Fourth was a model that incorporated both autoregression and encoding. Model four was the best-performing of these null models, so we used this as our benchmark.

For our SSM prediction metrics, we used the Cox-Snell ‘generalized’ *R*^2^ [75], which uses the likelihood ratio between competing models instead of the ratio of their residual variance:

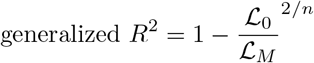

This metric is equivalent to the standard *R*^2^ for Gaussian models with independently and identically distributed residual covariance [75], but with the advantage that it reflects our model’s anisotropic, time-varying covariance from Kalman filtering.

#### 5.3 EEG-Optimized Recurrent Neural Networks

To compare SSMs to a more complex, non-linear model, we predicted the observed EEG (component) timeseries using a new set of RNNs implemented in PyTorch. We fit eight sets of RNNs, each yoked to the same number of free parameters as SSMs at each of the different levels of latent dimensionality. The inputs to this RNN were the previous timestep’s PC scores, and the same set of predictors as the SSM (formatted as constant inputs over each epoch, which fit better). These inputs were linearly projected into a set of hidden units, along with the previous hidden state, and then passed through a rectified linear unit (ReLU) activation function, using the standard PyTorch RNN implementation:

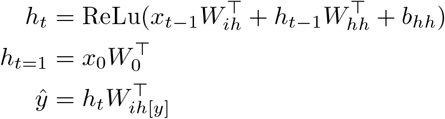

Note that for consistency with PyTorch, we use *h*_*t*_ to refer to the latent state (SSM: *x*_*t*_), and *x*_*t*_ to refer to the inputs (SSM: *u*_*t*_). *W*_*ih*[*y*]_ refers to the columns of *W*_*ih*_ used to encode the previous observations.

The PC scores on the next timestep were linearly decoded from the hidden state using the transpose of the observation embedding (a ‘tied’ parameterization; [76]). We found similar performance when we fit both the encoding and decoding layers, as this used more of our ‘free parameter budget’, trading off against the number of hidden units. The initial hidden state depended on the same inputs as the initial conditions of the SSM.

We fit 32 random initializations of this RNN to each participant and factor size. Each fitting session involved 5000 epochs of updating the parameters using backpropagation through time, using the mean square error of the next-timestep prediction as the loss function. Each batch contained all of the training data. Parameters were updated using the ‘AdamW’ learning rule (learning rate = .001, weight decay = .01; [77]). To evaluate each model’s performance, we took the best test-set loss across all epochs and initializations.

### 6 Dynamic Signatures of Task Control

#### 6.1 Additive state decomposition

To isolate the dynamics of task representations, we leveraged the powerful superposition property of linear systems. In this case, SSMs can be factorized into an additive set of subsystems for each input, a procedure known as ‘additive state decomposition’ [53]. We used this decomposition to model ‘task subsystems’, in which state dynamics are only caused by task inputs. We modeled task inputs for switch using the difference between ‘current task’ and ‘previous task’ predictors, and task inputs for repeat trials using their sum, providing separate task subsystems for switch and repeat trials. Within each subsystem, the state evolves according to:

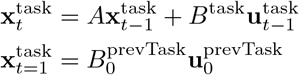

Since we contrast-coded the tasks (+1/-1), the dynamics for each task are symmetrical around the origin. For the initial conditions of this task subsystem, we used the ‘previous task’ component of the initial conditions 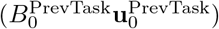, which is also symmetrical between switch and repeat trials for the same task.

To better equate task states across participants and datasets, we normalized the latent space using the univariate process noise (*T* = Diagonal(*W* ); *A*^*′*^ = *T* ^−1^*AT* ; *B*^*′*^ = *T* ^−1^*B*; *C*^*′*^ = *CT* ; see [42] *§*2.5).

#### 6.2 Switch Similarity

To characterize the similarity between task trajectories on switch and repeat trials, we measured the cross-lagged similarity between inferred task states for switch and repeat trials, similar to a temporal generalization analysis [55, 78]. In practice, we computed the cosine of the angle between task states across switch and repeat conditions at different cross-lags (i.e., computing the similarity between every timestep in the repeat subsystem to every timestep in the switch subsystem). We tested this similarity against zero at the group level using TFCE (Method 6.5).

#### 6.3 Initial Centrality

To understand whether task states converged to a common state during the ITI, we first measured the similarity between RNN hidden unit activations during the ITI. We averaged the hidden states during the ITI following each task, and then computed the (log) Euclidean distance between these average trajectories (i.e., the distances between the post-A states and the post-B states). Smaller values corresponded to states becoming more similar, and the slope of the line provided the (exponential) rate of convergence.

To understand whether RNNs and EEG converged to a ‘task neutral’ state during the ITI, we tested whether the terminal ITI state (i.e., at cue onset, the initial conditions of the SSM) was close to the midpoint of the states that were occupied during the trial. We developed a ‘midpoint score’ that quantified the relative Euclidean distance between the initial task state and each task state along the trajectory, which we computed for switch and repeat trials:

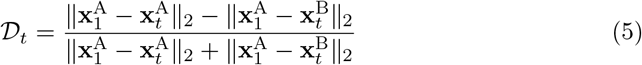

#### 6.4 Task Energy

To quantify the influence of task representations, we used energy-based methods from Lyapunov analysis. This control theoretic tool is used to quantify asymptotic system properties in methods like controllability analysis [43, 56, 79]. Controllability quantifies the strength of the input-state coupling. For a time-invariant system, the controllability Gramian (𝒢_∞_) defines the asymptotic state covariance that is attributable to a (white noise) input:

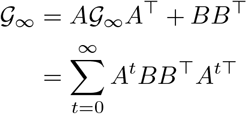

𝒢_∞_ can be computed by solving the corresponding Lyapunov equation. To capture the influence of a particular sequence of time-varying inputs, we used the recursive Lyapunov equation to provide a time-resolved task Gramian [58]:

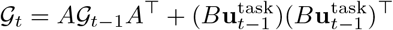

This task-dependent Gramian measures how much the system changes due to both immediate task inputs and the spread of task information through recurrence. At the initial condition, it has a value of zero. It is similar in spirit to a 2-norm on the task state [34], but additionally takes into account the cumulative state change (not just the distance from the origin). We summarized task energy using a standard metric of ‘average controllability’ [43], taking the trace of 𝒢 at each timestep, log transforming to reduce skew, and then taking the difference between Switch and Repeat conditions.

#### 6.5 Multiple comparisons corrections

To correct for multiple comparisons over time while accounting for temporal autocorrelation, we used threshold-free cluster enhancement throughout (TFCE; [80]) with a set of EEG-optimized parameters (H=2, C=1; from [81]; 1,000 permutations for temporal generalization and 10,000 permutations for traces).

### 7 EEG-RNN comparison

To evaluate the similarity between dynamic signatures between RNNs and humans, we first subsampled human task trajectories (100 or 125 timesteps) to match the duration of networks’ task trajectories (40 timesteps).

We averaged the outcome measures (Switch Similarity, Initial Centrality, or Task Energy) within RNNs and EEG, and then computed the similarity between RNNs and EEG. For Switch Similarity and Task Energy, we computed the similarity using the ‘congruence coefficient’, taking the cosine of the vectorized measures.

For Initial Centrality, we cared about the proximity to zero for a measure that was already normalized. For these reasons, we used an *R*^2^ similarity between EEG and RNNs:

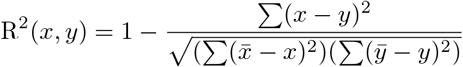

This metric gives a similarity value between 1 and . This metric can be negative if the distance between modalities is greater than the variance within modalities (recall that the modalities are not fit to each other).

To produce a bootstrap estimate of these metrics, we sampled with replacement 10,000 times within the RNN and EEG groups (e.g., drawing 512 RNNs and 30 EEG participants at random, with replacement). Within each bootstrap sample, we averaged our measures and computed our metrics as outlined above. We then computed the 95th percentiles of this bootstrap distribution.

To get an estimate of the difference between switch-trained RNNs, we sampled with replacement from (1) 1-Trial RNNs, (2) 2-Trial RNNs, and (3) EEG participants. We computed similarity metrics between 1-Trial RNNs and EEG, and between 2-Trial RNNs and EEG, and then we took the difference between these metrics. This approach provided a distribution of similarity differences, compared within the sample bootstrap sample of EEG participants. We expect EEG variability to be higher than RNNs due to noisy measurements and a smaller sample, and by accounting for EEG variability in this difference distribution, our difference measure could be more sensitive than looking for overlap in the confidence intervals for each metric.

### 8 EEG Task State Analyses

#### 8.1 EEG Task Trajectory Visualization

Using the decomposition and symmetry principles afforded by the linear system, we visualized the estimated latent trajectories of task representations between switch and repeat trials using group-level singular value decomposition (SVD).

To visualize the latent trajectories over time, we spatially concatenated participants’ switch and repeat task trajectories into a wide 2D ‘temporal’ matrix ((switch timesteps + repeat timesteps) *×* (factors *×*participants)). We then used SVD to extract separate scores for the switch and repeat subsystems. Since we only have two tasks, tasks were contrast coded (-1/1), which meant that the trajectories were symmetrical around an interpretable zero point (i.e., no task inputs). Despite this redundancy, we plotted the trajectories for all four conditions (i.e., task *×* switch) for illustrative purposes.

#### 8.2 EEG Performance Alignment

To confirm that the inferred task states were relevant for task performance, we tested whether trial-level brain states predicted upcoming reaction times.

First, we projected each participant’s task trajectory for switch and repeat subsystems back into the observation space 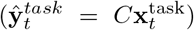. We computed the trial-averaged cosine similarity between the task state and the EEG response (mean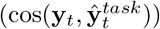). We included this ‘task score’ in a linear mixed-effects regression predicting log-transformed reaction times (MATLAB’s `fitlme`), alongside predictors accounting for the intercept, task, switch vs. repeat, and the task *×* switch interaction. We included maximal random intercepts and slopes, centered predictors within each participant, and confirmed that design matrices passed collinearity diagnostic tests. We tested whether group-level ‘task score’ estimates were significantly different from zero after the Satterthwaite correction for degrees of freedom. We found similar results when also controlling for the trial-averaged norm of the EEG and task state.

### 9 Software

1. Programming Languages
  a. julia (v1.11.2)
  b. python (v3.10.13)
  c. matlab (R2023b)
2. Software Packages
  a. PyTorch (v2.4.0, [64]) www.pytorch.org
  b. MatlabTFCE www.github.com/markallenthornton/MatlabTFCE
  c. ControlSystemIdentification.jl (v2.10.2) www.github.com/baggepinnen/ControlSystemIdentification.jl
  d. FieldTrip [82]
  e. EEGLAB (v2024.2; [83])
3. Visualization
  a. Scientific Color Maps [84]

The author(s) are pleased to acknowledge that the work reported on in this paper was substantially performed using the Princeton Research Computing resources at Princeton University, which is a consortium of groups led by the Princeton Institute for Computational Science and Engineering (PICSciE) and the Office of Information Technology’s Research Computing.

## Supplementary Figures

**Fig. 1:**
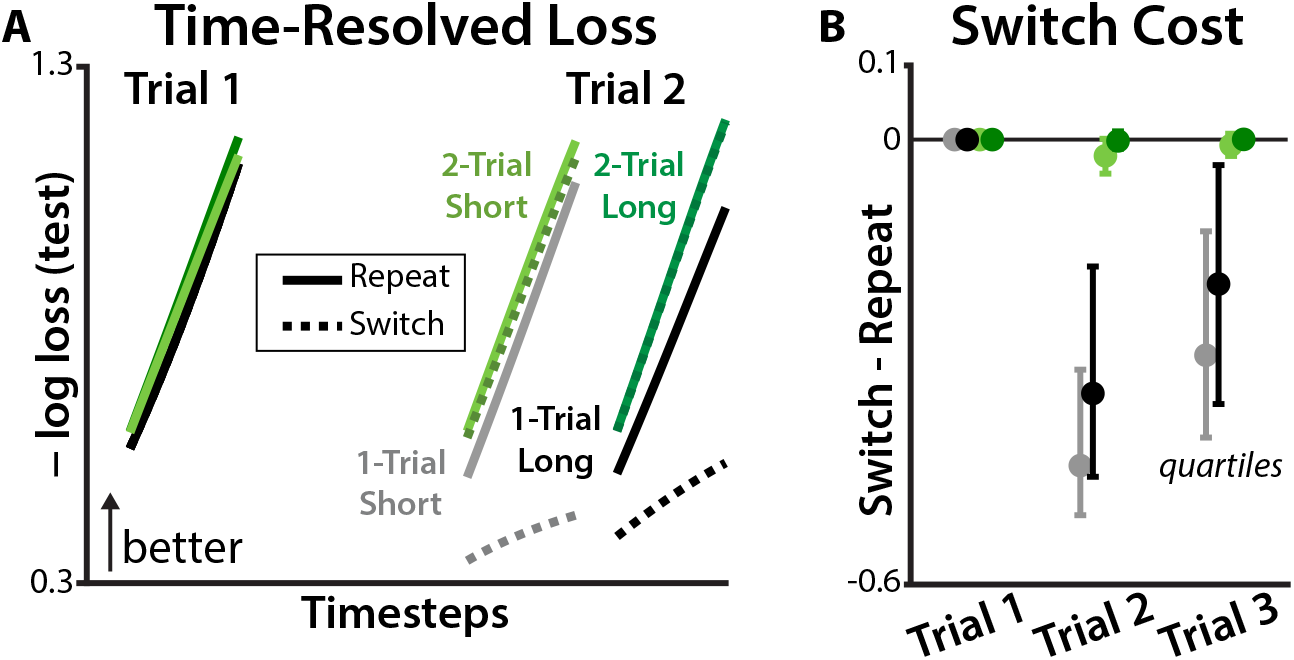
Differences in switch costs across RNNs. **A)** Time-resolved test loss for each RNN condition, plotted for two trials that either switched task (solid line) or repeated tasks (dashed line). Loss is averaged over networks within each RNN condition; SEM not show because they were similar to or smaller than the line width. Note that the second trial for Long-ITI RNNs are delayed due to their longer ITI. **B)** Difference in the test loss switch and repeat trials, plotted for each RNN condition on 3-trial sequences. Error bars reflect quartiles of the between-RNN distribution.

**Fig. 2:**
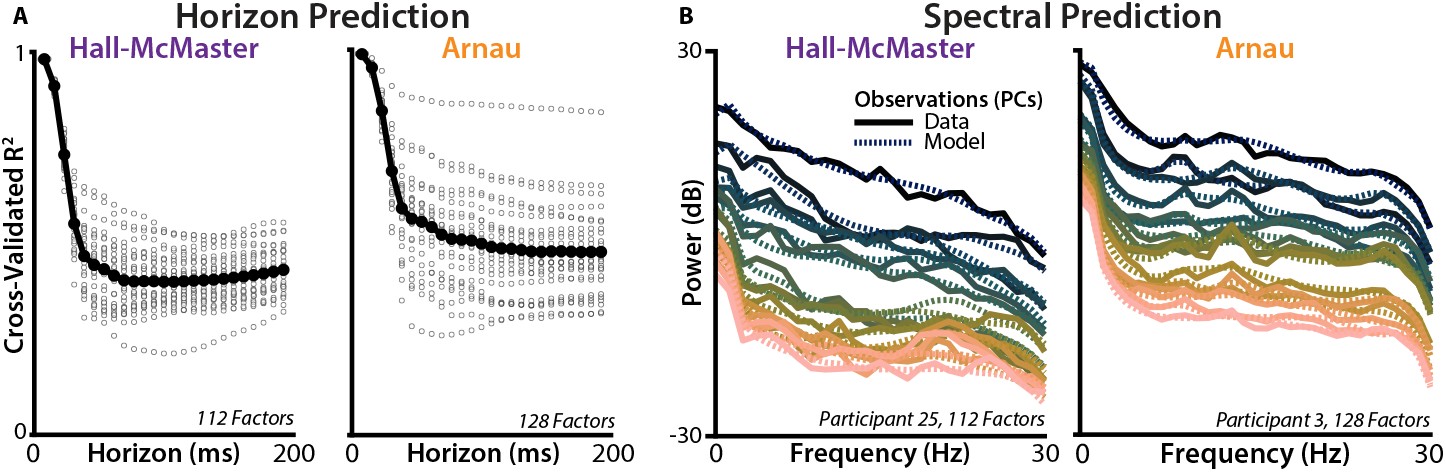
Long-range predictive accuracy of SSMs fits to EEG. **A)** Predictive accuracy of the model when forecasting at different timesteps into the future. Gray dots indicate individual participants. **B)** Similarity in the power spectrum for test-set observations (solid lines) and model simulations (dashed lines), shown for example participants. Rather than using Kalman filtered predictions, these predictions were generated from 100 posterior samples per participant (i.e., feedforward roll-outs). Power spectra are plotted for each observation dimension (i.e., principal component), aggregated across stimulated epochs using EEGLab’s `spectopo` function. Across participants, SSMs accurately captured the empirical power spectra (Hall-McMaster: 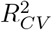 95% CI [0.97, 0.98]; Arnau: 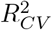 95% CI [0.99, 0.99]). We estimated the the noise ceiling for power spectrum similarity using the similarity between the empirical training-set spectrum and the empirical test-set spectrum, finding that SSMs captured most of the reliable signal (Hall-McMaster: noise-normalized 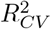 95% CI [.99, .99]; Arnau: noise-normalized 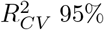 CI [0.99, 1.0]). There was still high overlap between model and data after mean-centering and linearly detrending the spectra within each principal component (Hall-McMaster: 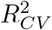 95% CI [0.79, 0.88], noise-normalized 95% CI [0.92, 0.94]; Arnau: 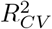 95% CI [0.87, 0.93], noise-normalized 95% CI [0.89, 0.97]).

**Fig. 3:**
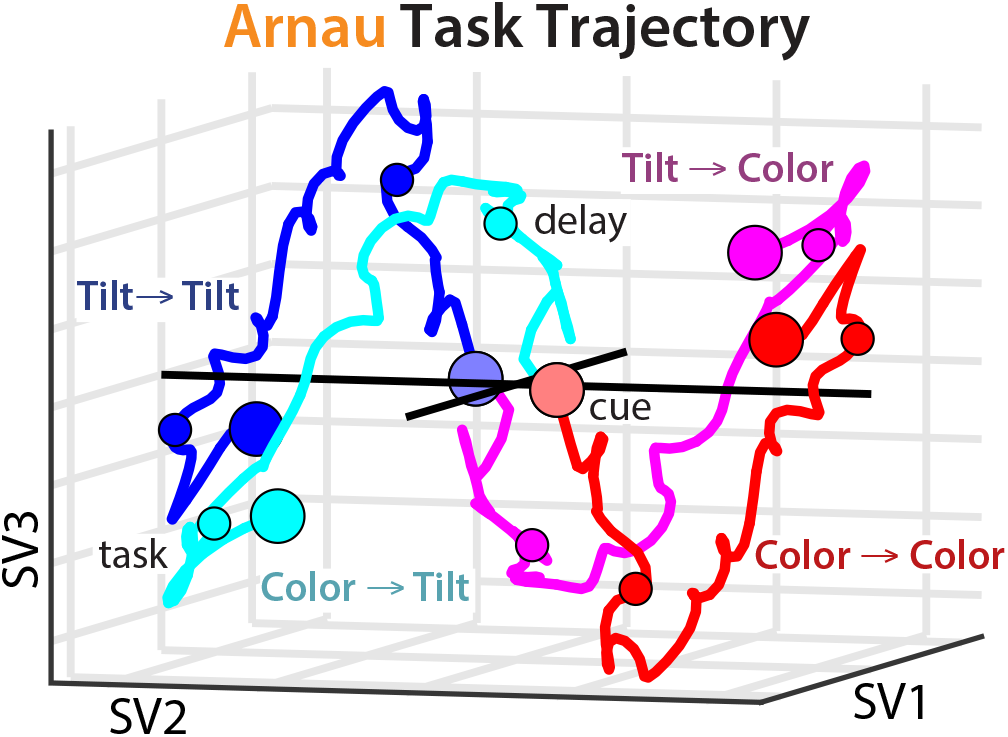
Latent task trajectories from SSMs fit to Arnau. Low-dimensional embedding of the expected task trajectories, split by switch and repeat conditions (SV: singular vector). Task is contrast-coded, so task trajectories within the same switch condition are symmetrical. Shown for Arnau, see Fig. 4C for Hall-McMaster.

**Fig. 4:**
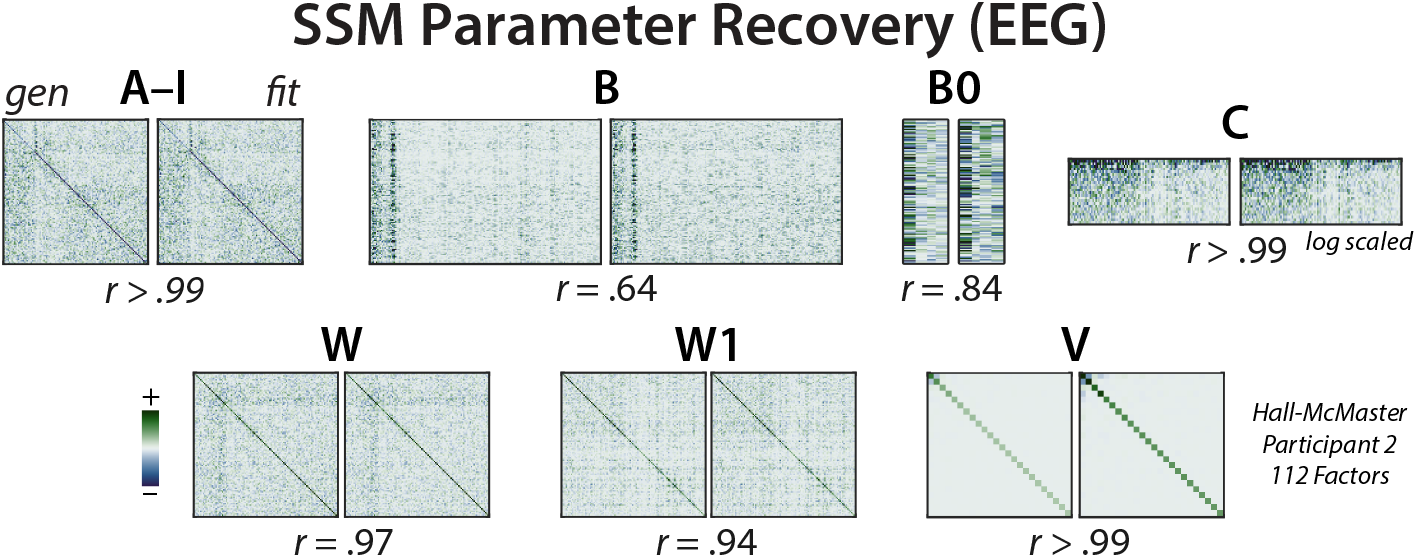
SSM parameter recovery in EEG. Generating and fit parameters for an example participant in Hall-McMaster. Parameters were linearly aligned to account for degeneracy in the latent system Method 4.4.

## Notes

https://github.com/harrisonritz/StateSpaceAnalysis.jl

### Competing Interest Statement

The authors have declared no competing interest.

### Summary of Updates

Major structural revision of the paper, but data, analyses, and interpretation are consistent with previously uploaded version.

## References

[1] Monsell, S. Task switching. Trends Cogn. Sci. 7, 134–140 (2003).

[2] Kiesel, A. et al. Control and interference in task switching—a review. Psychol. Bull. 136, 849–874 (2010).

[3] Vandierendonck, A., Liefooghe, B. & Verbruggen, F. Task switching: interplay of reconfiguration and interference control. Psychol. Bull. 136, 601–626 (2010).

[4] Musslick, S. & Cohen, J. D. Rationalizing constraints on the capacity for cognitive control. Trends Cogn. Sci. 0 (2021).

[5] Friedman, N. P. & Miyake, A. Unity and diversity of executive functions: Indi-vidual differences as a window on cognitive structure. Cortex 86, 186–204 (2017).

[6] Cepeda, N. J., Kramer, A. F. & Gonzalez de Sather, J.C.M. Changes in executive control across the life span: Examination of task-switching performance. Dev. Psychol. 37, 715–730 (2001).

[7] Steyvers, M., Hawkins, G. E., Karayanidis, F. & Brown, S. D. A large-scale analysis of task switching practice effects across the lifespan. Proc. Natl. Acad. Sci. U. S. A. (2019).

[8] Millan, M. J. et al. Cognitive dysfunction in psychiatric disorders: characteristics, causes and the quest for improved therapy. Nat. Rev. Drug Discov. 11, 141–168 (2012).

[9] Snyder, H. R. Major depressive disorder is associated with broad impairments on neuropsychological measures of executive function: a meta-analysis and review. Psychol. Bull. 139, 81–132 (2013).

[10] Goschke, T. Intentional reconfiguration and involuntary persistence in task-set switching. Control of cognitive processes 331–355 (2000).

[11] Meiran, N. Modeling cognitive control in task-switching. Psychol. Res. 63, 234– 249 (2000).

[12] Rogers, R. D. & Monsell, S. Costs of a predictible switch between simple cognitive tasks. J. Exp. Psychol. Gen. 124, 207 (1995).

[13] Monsell, S. in Task set regulation (ed. Egner, T.) The Wiley Handbook of Cognitive Control, Vol. 115 29–49 (John Wiley & Sons, Ltd, Chichester, UK, 2017).

[14] Allport, A., Styles, E. A. & Hsieh, S. L. in Shifting intentional set - exploring the dynamic control of tasks 421–452 (MIT Press, London, England, 1994).

[15] Wylie, G. & Allport, A. Task switching and the measurement of “switch costs”. Psychol. Res. 63, 212–233 (2000).

[16] Meiran, N. Reconfiguration of processing mode prior to task performance. J. Exp. Psychol. Learn. Mem. Cogn. 22, 1423–1442 (1996).

[17] Yeung, N. & Monsell, S. Switching between tasks of unequal familiarity: the role of stimulus-attribute and response-set selection. J. Exp. Psychol. Hum. Percept. Perform. 29, 455–469 (2003).

[18] Schmitz, F. & Voss, A. Decomposing task-switching costs with the diffusion model. J. Exp. Psychol. Hum. Percept. Perform. 38, 222–250 (2012).

[19] Yeung, N., Nystrom, L. E., Aronson, J. A. & Cohen, J. D. Between-task competition and cognitive control in task switching. J. Neurosci. 26, 1429–1438 (2006).

[20] Loose, L. S., Wisniewski, D., Rusconi, M., Goschke, T. & Haynes, J.-D. Switchindependent task representations in frontal and parietal cortex. J. Neurosci. 37, 8033–8042 (2017).

[21] Qiao, L., Zhang, L., Chen, A. & Egner, T. Dynamic trial-by-trial recoding of taskset representations in the frontoparietal cortex mediates behavioral flexibility. J. Neurosci. 37, 11037–11050 (2017).

[22] Gilbert, S. J. & Shallice, T. Task switching: a PDP model. Cogn. Psychol. 44, 297–337 (2002).

[23] Ardid, S. & Wang, X.-J. A tweaking principle for executive control: neuronal circuit mechanism for rule-based task switching and conflict resolution. J. Neurosci. 33, 19504–19517 (2013).

[24] Musslick, S., Jang, S. J., Shvartsman, M., Shenhav, A. & Cohen, J. D. Constraints associated with cognitive control and the stability-flexibility dilemma (shenhavlab.org, 2018).

[25] Ueltzhöffer, K., Armbruster-Genç, D. J. N. & Fiebach, C. J. Stochastic dynamics underlying cognitive stability and flexibility. PLoS Comput. Biol. 11, e1004331 (2015).

[26] Jaffe, P. I., Poldrack, R. A., Schafer, R. J. & Bissett, P. G. Modelling human behaviour in cognitive tasks with latent dynamical systems. Nature Human Behaviour 1–15 (2023).

[27] Cho, K. et al. Learning phrase representations using RNN encoder-decoder for statistical machine translation. arXiv [cs.CL] (2014).

[28] Siegel, M., Donner, T. H. & Engel, A. K. Spectral fingerprints of large-scale neuronal interactions. Nat. Rev. Neurosci. 13, 121–134 (2012).

[29] Mante, V., Sussillo, D., Shenoy, K. V. & Newsome, W. T. Context-dependent computation by recurrent dynamics in prefrontal cortex. Nature 503, 78–84 (2013).

[30] Cohen, J. D., Dunbar, K. & McClelland, J. L. On the control of automatic processes: a parallel distributed processing account of the stroop effect. Psychol. Rev. 97, 332–361 (1990).

[31] Braver, T. S. & Cohen, J. D. in Chapter 19 dopamine, cognitive control, and schizophrenia: the gating model , Vol. 121 of Progress in brain research 327–349 (Elsevier, 1999).

[32] O’Reilly, R. C. & Frank, M. J. Making working memory work: a computational model of learning in the prefrontal cortex and basal ganglia. Neural Comput. 18, 283–328 (2006).

[33] Sussillo, D. & Barak, O. Opening the black box: low-dimensional dynamics in high-dimensional recurrent neural networks. Neural Comput. 25, 626–649 (2013).

[34] Soldado-Magraner, J., Mante, V. & Sahani, M. Inferring context-dependent computations through linear approximations of prefrontal cortex dynamics. Sci. Adv. 10, eadl4743 (2024).

[35] Gurnani, H., Liu, W. & Brunton, B. W. Feedback control of recurrent dynamics constrains learning timescales during motor adaptation. bioRxiv 2024.05.24.595772 (2024).

[36] Brown, E. N., Frank, L. M., Tang, D., Quirk, M. C. & Wilson, M. A. A statistical paradigm for neural spike train decoding applied to position prediction from ensemble firing patterns of rat hippocampal place cells. J. Neurosci. 18, 7411– 7425 (1998).

[37] Smith, A. C. & Brown, E. N. Estimating a state-space model from point process observations. Neural Comput. 15, 965–991 (2003).

[38] Macke, J. H. et al. Empirical models of spiking in neural populations (2011).

[39] LaFosse, P. K. et al. Single-cell optogenetics reveals attenuation-by-suppression in visual cortical neurons. bioRxivorg 2023.09.13.557650 (2024).

[40] Nozari, E. et al. Macroscopic resting-state brain dynamics are best described by linear models. Nat. Biomed. Eng. 1–17 (2023).

[41] Jha, A., Gupta, D., Brody, C. D. & Pillow, J. W. Disentangling the roles of distinct cell classes with cell-type dynamical systems. bioRxiv 2024.07.08.602520 (2024).

[42] Williams, R. L. & Lawrence, D. A. Linear state-space control systems (John Wiley & Sons, Nashville, TN, 2007).

[43] Tang, E. & Bassett, D. S. Colloquium: Control of dynamics in brain networks. Rev. Mod. Phys. (2018).

[44] Linderman, S., Nichols, A., Blei, D., Zimmer, M. & Paninski, L. Hierarchical recurrent state space models reveal discrete and continuous dynamics of neural activity in C. elegans. bioRxiv 621540 (2019).

[45] Gohil, C. et al. Dynamic network analysis of electrophysiological task data. Imaging Neuroscience 2, 1–19 (2024).

[46] Misra, J. & Pessoa, L. Brain dynamics and spatiotemporal trajectories during threat processing (2025).

[47] Hess, F., Monfared, Z., Brenner, M. & Durstewitz, D. Generalized teacher forcing for learning chaotic dynamics, 13017–13049 (PMLR, 2023).

[48] Pals, M., Sağtekin, A. E., Pei, F., Gloeckler, M. & Macke, J. H. Inferring stochastic low-rank recurrent neural networks from neural data. arXiv [cs.LG] (2024).

[49] Ghahramani, Z., Rey, G. & Hinton, E. Parameter estimation for linear dynamical systems. Techinical Report (1996).

[50] Murphy, K. P. Probabilistic Machine Learning: Advanced Topics (MIT Press, London, England, 2023).

[51] Ritz, H. harrisonritz/StateSpaceAnalysis.jl: v0.2.0 (2024).

[52] Brunton, S. L., Brunton, B. W., Proctor, J. L., Kaiser, E. & Kutz, J. N. Chaos as an intermittently forced linear system. Nat. Commun. 8, 19 (2017).

[53] Staffans, O. Encyclopedia of mathematics and its applications: Well-posed linear systems series number 103 (Cambridge University Press, Cambridge, England, 2009).

[54] King, J. A., Korb, F. M., von Cramon, D. Y. & Ullsperger, M. Post-error behavioral adjustments are facilitated by activation and suppression of task-relevant and task-irrelevant information processing. J. Neurosci. 30, 12759–12769 (2010).

[55] Luyckx, F., Nili, H., Spitzer, B. & Summerfield, C. Neural structure mapping in human probabilistic reward learning. Elife 8 (2019).

[56] Kao, T.-C. & Hennequin, G. Neuroscience out of control: control-theoretic perspectives on neural circuit dynamics. Curr. Opin. Neurobiol. 58, 122–129 (2019).

[57] Holroyd, C. B. The controllosphere: The neural origin of cognitive effort. Psychol. Rev. (2024).

[58] Klett, C., Abate, M., Yoon, Y., Coogan, S. & Feron, E. Bounding the state covariance matrix for switched linear systems with noise, 2876–2881 (IEEE, 2020).

[59] Hall-McMaster, S., Muhle-Karbe, P. S., Myers, N. E. & Stokes, M. G. Reward boosts neural coding of task rules to optimize cognitive flexibility. J. Neurosci. 39, 8549–8561 (2019).

[60] Arnau, S., Liegel, N. & Wascher, E. Frontal midline theta power during the cue-target-interval reflects increased cognitive effort in rewarded task-switching. Cortex 180, 94–110 (2024).

[61] Athalye, V. R., Carmena, J. M. & Costa, R. M. Neural reinforcement: reentering and refining neural dynamics leading to desirable outcomes. Curr. Opin. Neurobiol. 60, 145–154 (2019).

[62] Ritz, H., Leng, X. & Shenhav, A. Cognitive control as a multivariate optimization problem. J. Cogn. Neurosci. 34, 569–591 (2022).

[63] Fagot, C. & Pashler, H. Making two responses to a single object: implications for the central attentional bottleneck. J. Exp. Psychol. Hum. Percept. Perform. 18, 1058–1079 (1992).

[64] Paszke, A. et al. PyTorch: An imperative style, high-performance deep learning library. arXiv [cs.LG] (2019).

[65] Ehrlich, D. B. & Murray, J. D. Geometry of neural computation unifies working memory and planning. Proc. Natl. Acad. Sci. U. S. A. 119, e2115610119 (2022).

[66] Langdon, C. & Engel, T. A. Latent circuit inference from heterogeneous neural responses during cognitive tasks. Nat. Neurosci. 1–11 (2025).

[67] Linderman, S. W. et al. Dynamax: A python package for probabilistic state space modeling with JAX. J. Open Source Softw. 10, 7069 (2025).

[68] Holmes, e., Elizabeth, Ward, j., Eric & Wills, K. MARSS: Multivariate autoregressive state-space models for analyzing time-series data. R J. 4, 11 (2012).

[69] Sarkka, S. Institute of mathematical statistics textbooks: Bayesian filtering and smoothing series number 3 (Cambridge University Press, Cambridge, England, 2013).

[70] Smith, G., de Freitas, J., Robinson, T. & Niranjan, M. Speech modelling using subspace and EM techniques. Advances in Neural Information Processing Systems 12 (1999).

[71] Stone, I. R., Sagiv, Y., Park, I. M. & Pillow, J. W. Spectral learning of bernoulli linear dynamical systems models. arXiv [stat.ML] (2023).

[72] Larimore, W. E. Canonical variate analysis in identification, filtering, and adaptive control, 596–604 vol.2 (IEEE, 1990).

[73] Stephan, K. E., Penny, W. D., Daunizeau, J., Moran, R. J. & Friston, K. J. Bayesian model selection for group studies. Neuroimage 46, 1004–1017 (2009).

[74] Ehinger, B. V. & Dimigen, O. Unfold: an integrated toolbox for overlap correction, non-linear modeling, and regression-based EEG analysis. PeerJ 7, e7838 (2019).

[75] Cox, D. R. & Wermuth, N. A comment on the coefficient of determination for binary responses. Am. Stat. 46, 1–4 (1992).

[76] Hinton, G. E. & Salakhutdinov, R. R. Reducing the dimensionality of data with neural networks. Science 313, 504–507 (2006).

[77] Loshchilov, I. & Hutter, F. Decoupled weight decay regularization (2017).

[78] King, J.-R. & Dehaene, S. Characterizing the dynamics of mental representations: the temporal generalization method. Trends Cogn. Sci. 18, 203–210 (2014).

[79] Kalman, R. E. in Lectures on controllability and observability 1–149 (Springer Berlin Heidelberg, Berlin, Heidelberg, 2010).

[80] Smith, S. M. & Nichols, T. E. Threshold-free cluster enhancement: addressing problems of smoothing, threshold dependence and localisation in cluster inference. Neuroimage 44, 83–98 (2009).

[81] Mensen, A. & Khatami, R. Advanced EEG analysis using threshold-free clusterenhancement and non-parametric statistics. Neuroimage 67, 111–118 (2013).

[82] Oostenveld, R., Fries, P., Maris, E. & Schoffelen, J.-M. FieldTrip: Open source software for advanced analysis of MEG, EEG, and invasive electrophysiological data. Comput. Intell. Neurosci. 2011, 156869 (2011).

[83] Delorme, A. & Makeig, S. EEGLAB: an open source toolbox for analysis of singletrial EEG dynamics including independent component analysis. J. Neurosci. Methods 134, 9–21 (2004).

[84] Crameri, F., Shephard, G. E. & Heron, P. J. The misuse of colour in science communication. Nat. Commun. 11, 5444 (2020).

